# Comparing the potential of microglia preparations generated using different human iPSC-based differentiation methods to model microglia-mediated mechanisms of amyotrophic lateral sclerosis pathophysiology

**DOI:** 10.1101/2022.03.11.483851

**Authors:** Ye Man Tang, Nisha S. Pulimood, Stefano Stifani

**Affiliations:** Department of Neurology and Neurosurgery, Montreal Neurological Institute-Hospital, McGill University, Montreal, Quebec, H3A 2B4 Canada

**Keywords:** amyotrophic lateral sclerosis, heterogeneity, human iPSCs, microglia, RNA sequencing

## Abstract

Microglia are the resident immune cells of nervous system. In healthy conditions, microglia actively patrol neural tissues in a homeostatic state, which can rapidly change to an activated state in response to local injury/disease. Dysregulated microglia activation is a hallmark of disorders and diseases of the nervous system, including motor neuron diseases such as amyotrophic lateral sclerosis (ALS). The elucidation of the roles of microglia in human biology and disease has recently benefitted from the development of human induced pluripotent stem cell (iPSC)-based approaches to generate microglia-like cells. Microglia represent a heterogeneous group of cells with spatial diversity in both health and disease. This situation poses a considerable challenge along the path towards establishing the most pathologically-relevant human iPSC-derived microglia preparations to investigate the complex roles of microglia in ALS and other neurological diseases. The success of these approaches must account for microglia diversity in different regions of the brain and spinal cord. In this study, we compared the transcriptomes of human iPSC-derived microglia generated using different methods to determine whether or not separate strategies can be used to generate microglia with distinct transcriptional signatures *in vitro*. We show that microglia derived using two different methods display distinct *in vitro* maturation characteristics. We also reveal that different derivation methods give rise to preparations comprising human microglia with distinct transcriptomic signatures resembling the gene profiles of specific microglia subpopulations *in vivo*. These findings suggest that a careful, and coordinated, implementation of multiple microglia differentiation methods from human iPSCs can be an effective approach towards the goal of generating multiple microglia subtypes that will offer enhanced model systems to account for microglia heterogeneity *in vivo*. Spatially-defined human iPSC-derived microglia would represent an enhanced tool to study the multiple levels of involvement of microglia in mechanisms of motor neuron degeneration in ALS, as well as other neurological diseases and disorders.

## 1. Introduction

Microglia are the resident myeloid cells of the adult nervous system. They mediate innate immunity by sensing and controlling changes in the microenvironment caused by either external pathogens or local pathologies. In healthy conditions, microglia patrol neural tissues in a homeostatic (‘resting’) state, which can rapidly change to an ‘activated’ state in response to injury/disease. Activated microglia are the primary phagocytic cells in the nervous system. They can engulf external pathogens and cellular debris resulting from injury or disease. Activated microglia also release inflammatory cytokines that can promote astrocyte activation and be toxic to neighbouring neurons. The balanced contributions of resting and activated microglia mediate both beneficial (protective) and detrimental (inflammatory) effects on the surrounding cells, depending on the specific context (reviewed by Prinz *et al*., 2019; Miron & Priller, 2020; Mendes & Majewska, 2021; Spiteri *et al*., 2022).

The presence of activated microglia is a common event in virtually all neurological diseases and disorders. For instance, hallmarks of microglia activation are a feature of *post-mortem* tissues from amyotrophic lateral sclerosis (ALS) patients (Yasojima *et al*., 2001; Henkel *et al*., 2004; Malaspina & de Belleroche, 2004; Lederer *et al*., 2007), as well as spinal cord and cranial motor nuclei of ALS transgenic mouse models (Fendrick *et al*., 2007; Henkel *et al*., 2009; Kassa *et al*., 2009; Graber *et al*., 2010). ALS is an incurable motor neuron disease characterized by the progressive death of motor neurons in the cerebral cortex (‘upper motor neurons’), brain stem and spinal cord (‘lower motor neurons’) (reviewed by Grad *et al*., 2017; van Es *et al*., 2017; Mejzini *et al*., 2019). Intercellular communication between motor neurons and microglia is proposed to play important and dynamic roles during the degeneration of motor circuits in ALS. Microglia are believed to initially protect motor neurons during early phases of the disease by performing scavenging, anti-inflammatory, and neuroprotective functions. At more advanced stages, however, with rising oxidative stress and injury within neurons, microglia can contribute to motor neuron death by switching to pro-inflammatory and neurotoxic phenotypes (reviewed by Geloso *et al*., 2017; Beers & Appel, 2019; McCauley & Baloh, 2019; Clarke & Patani, 2020).

The study of the mechanisms underlying the roles of microglia in ALS pathophysiology, as well as other neurodegenerative conditions such as Parkinson’s disease and Alzheimer’s disease, has benefitted from the availability of animal experimental model systems (reviewed by Thonhoff *et al*., 2018; Beers & Appel, 2019; Karanfilian *et al*., 2020; Liu *et al*., 2021). However, numerous species-specific differences in gene expression exist between murine and human microglia. Moreover, several microglial genes exhibiting differences among species are often implicated in human neurodegenerative diseases (Szulzewsky *et al*., 2016; Masuda *et al*., 2019; Patir *et al*., 2019; Healy *et al*., 2020). These observations highlight the need to combine studies in animal models with investigations using human microglia-based experimental systems. Recent advances in induced pluripotent stem cell (iPSC)-based approaches have addressed this need by providing efficient and reliable approaches to generate physiologically-relevant human microglia-like cells (e.g. Muffat *et al*., 2016; Abud *et al*., 2017; Douvaras *et al*., 2017; Pandya *et al*., 2017; McQuade *et al*., 2018; to cite a few). This remarkable progress offers previously unavailable experimental opportunities to study human microglia biology and model microglia pathophysiological mechanisms *in vitro*.

Recent studies have provided growing evidence that microglia represent a heterogeneous group of cells comprising region-specific subpopulations with distinct biological characteristics (Grabert *et al*., 2016; DePaula-Silva *et al*., 2019; Li *et al*., 2019; Masuda *et al*., 2019; Stratoulias *et al*., 2019; van der Poel *et al*., 2019; Masuda *et al*., 2020). This cellular heterogeneity poses a challenge to research strategies involving microglia-like cells derived from human iPSCs (hereafter referred to as ‘human iPSC-derived microglia’). It is reasonable to postulate that the success of these investigations will depend on whether or not human iPSC-derived microglia preparations will comprise those specific subtypes that are more relevant to the biological questions under study. A case in point: neuroinflammatory responses mediated by microglia in ALS are heterogeneous, resulting in the presence of distinct subsets of activated microglia that are believed to interact with disease-specific motor neurons in regionally-defined patterns (Dachet *et al*., 2019; Maniatis *et al*., 2019; Cipollina *et al*., 2020; Liu *et al*., 2021). More generally, the field of nervous system disease modeling using human iPSCs is increasingly recognizing the importance of including the *in vivo* heterogeneity of the induced cells among the factors that need to be considered when designing ideal, disease-relevant cellular assays (Hedegaard *et al*., 2020; Stifani, 2021; Giacomelli *et al*., 2022).

In this study, we provide evidence that human iPSC-derived microglia preparations generated using different methods comprise microglia with distinct transcriptional signatures *in vitro* resembling gene profiles of different microglia subpopulations *in vivo*. These observations suggest that the harmonised use of multiple human microglia derivation methods should be implemented when modeling ALS, as well as other neurodegenerative diseases, in order to generate the most disease-relevant repertoire of microglia subgroups needed to model pathophysiological mechanisms involving regionally-specialized microglia subtypes.

## 2. Materials and Methods

### 2.1. Human induced pluripotent stem cellsv

Human iPSC line NCRM-1 was obtained from the National Institutes of Health Stem Cell Resource (Bethesda, MD, USA). Human iPSC lines CS52iALS-C9n6.ISOxx and CS29iALS-C9n1.ISOxx, corresponding to corrected, isogenic lines derived from two distinct parental lines generated from ALS patients with *C9orf72* gene mutations, were obtained from Cedars-Sinai (Los Angeles, CA, USA) (Thiry *et al*., 2022). Maintenance and quality control of iPSCs was conducted as described previously (Soubannier *et al*., 2022).

### 2.2. Derivation of microglia-like cells from human iPSCs

Derivation of microglia from human iPSCs (n=3 lines) was performed as described in two separate protocols published by Fossati and coworkers (Douvaras *et al*., 2017) and Blurton-Jones and colleagues (McQuade *et al*., 2018). The following modifications were made to the Douvaras *et al* method. Human iPSCs were plated onto 6-well dishes coated with Matrigel (Thermo-Fisher Scientific; Waltham, MA, USA; Cat. No. 08-774-552) in mTeSR1 medium (STEMCELL Technologies; Vancouver, BC, Canada; Cat. No. 85850) containing 10 µM ROCK inhibitor (compound Y-27632 2HCl; Selleck Chemicals; Cat. No. S1049) for 24 hours (this was considered as day 0 *in vitro*). Starting on the next day, the culture medium was changed to Essential 6 Medium (ThermoFisher Scientific; Cat. No. A1516401) containing 80 ng/ml BMP4 and cells were cultured for 2-4 days until they reached near confluence. All subsequent steps were as described in the original protocol (Douvaras *et al*., 2017), except that microglial progenitors were routinely collected as floating cells, without need for fluorescence activated cell sorting or magnetic bead separation, followed by culture for further differentiation in SF-Microglia Medium (RPMI + IL-34 + GM-CSF).

### 2.3. Characterization of human iPSC-derived microglia by immunocytochemistry

Induced human microglia were analyzed by immunocytochemistry, which was performed as described previously (Methot *et al*., 2018), using the following primary antibodies: rabbit anti-IBA1 (1/1,000; FUJIFILM Wako Chemicals; Richmond, VA, USA; Cat. No.019-19741; rabbit anti-P2Y12R (1/150; Alomone Labs; Jerusalem, Israel; Cat. No. APR-020). The secondary antibody was a donkey anti-rabbit IgG conjugated to Alexa Fluor 594 (1/1,000; Invitrogen; Burlington, ON, Canada; Cat No. A-21207; RRID, AB_141637). Images were acquired using a Zeiss Axio Imager M1 microscope connected to an AxioCam 503 camera, using ZEN software (Zeiss Canada Ltd., Toronto, ON, Canada).

### 2.4. RNA sequencing and acquisition of read counts

Total RNA was isolated for RNA sequencing (RNAseq) from cell pellets via sequential treatment with TRIzol Reagent (ThermoFisher Scientific; Cat. No. 15596026) and PureLink RNA Micro Scale Kit (ThermoFisher Scientific; Cat. No. 12183-016) following the instructions provided by the manufacturer. RNA samples were submitted for Illumina next-generation sequencing to the Genomics platform at the Institute for Research in Immunology and Cancer, Montreal, QC, Canada (https://www.iric.ca/en/research/platforms-and-infrastructures/genomics). Adaptor sequences and low-quality bases in the resulting FASTQ files were trimmed using Trimmomatic version 0.35 (Bolger *et al*., 2014) and genome alignments were conducted using STAR version 2.5 1b (Dobin *et al*., 2013). The sequences were aligned to the human genome version GRCh38, with gene annotations from Gencode v29 based on Ensemble release 94. As part of quality control, the sequences were aligned to several different genomes to verify that there was no sample contamination. Raw read-counts were obtained directly from STAR and reads in transcripts per million (TPM) format were computed using RSEM (Li & Dewey, 2011).

### 2.5. In silico analysis

Heatmaps were used to visualize expression of specific gene sets across multiple different samples. To generate heatmaps, read counts were converted into log2-counts-per-million (logCPM) values with the “cpm” function of the edgeR package in R. This function adds a low “prior count” value to avoid taking the log of zero and also normalizes that data by the different library sizes of the samples. Accordingly, negative values represent very low gene expression values, usually below 5. Hierarchical clustering of rows was applied to the heatmaps in order to group genes with correlated expression patterns. Since logCPM values are displayed directly without additional Z-score scaling by row, color comparisons can be made across samples for each gene as well as between genes for any given sample.

Differential gene expression analysis was conducted on the raw read-count matrix using an edgeR (version 4.0) pipeline as previously described (Chen *et al*., 2016). Briefly, we first filtered the genes such that only those with a robust expression level were retained. Next, the differing RNA composition/library size of each sample was normalized by a scaling factor of the trimmed mean of M-values (TMM) between each pair of samples. The known sources of variability in the studied samples were the microglia differentiation protocol and batch, both of which were incorporated into the experimental design used for differential expression analysis. The “glmQLFit” and “glmQLFTest” functions of edgeR were implemented to obtain differentially expressed genes (DEGs) using the quasi-likelihood (QL) method. Genes were considered statistically significant if the adjusted p-value (false discovery rate; FDR) was < 0.05, with no fold-change cutoff applied to restrict the number of DEGs identified.

The GeneOverlap package in R was used to reveal statistically significant overlaps between gene sets in comparison to the genomic background. We used the DEGs upregulated in microglia derived using either protocol and compared them with previously published gene signatures for different microglial subtypes. All the gene sets used in this analysis are listed in Supplementary Table 1 available in Supplementary Materials. Fisher’s exact test was used to generate the overlapping p-value, with the null hypothesis that the odds ratio ≤ 1. Odds ratio is a statistic that quantifies the strength of association between two events (in this case, gene sets); an odds ratio ≥ 1 means there is a correlation between the two gene sets.

Gene set enrichment analysis (GSEA) was conducted on the DEGs using the Functional Annotation Tool of DAVID Bioinformatics Resources 6.8 (Huang *et al*., 2009a; 2009b). Gene Ontology (GO) terms and Kyoto Encyclopedia of Genes and Genomes (KEGG) pathways enriched in the DEGs were computed separately for upregulated and downregulated genes, as this increases the statistical power to identify pathways/terms pertinent to the phenotype of interest (Hong *et al*., 2013). For each GO category (for example, Biological Process or Molecular Function), the “direct” function in DAVID was used to restrict the results to those directly annotated by the source database and to avoid repetitive and generic parent terms. For the KEGG pathways, only the non-disease-specific pathways are shown.

### 2.6. Open source data

RNAseq data (raw read counts) for iPSC lines were obtained from the LINCS Data Portal (LSC-1002 and LSC-1004 from Dataset LDS-1355) (http://lincsportal.ccs.miami.edu/datasets/). LSC-1002 and LSC-1004 correspond to iPSC lines CS14i-CTR-n6 and CS25iCTR-18n2, whose source provider was Cedar Sinai Stem Cell Core Laboratory. RNAseq data for the microglia line C20 were obtained from the Digital Expression Explorer 2 (DEE2) repository (Accession# SRR3546647) (http://dee2.io/). All these raw data were combined with the RNAseq data from our microglia and analyzed using a single pipeline. Additional metadata for all open source RNAseq data are provided in Supplementary Materials (Supplementary Table 2).

## 3. Results

### 3.1. Characterization of human iPSC-derived microglia generated using different protocols

The *in vivo* heterogeneity of microglia is seldom considered when human iPSCs are used to derive microglial cells to study their roles in (patho)biological mechanisms in health and disease. This omission has repercussions when human iPSC-derived microglia with no defined regional characteristics are used to model neurodegenerative diseases that affect specific neuronal cell types in specific regions of the central nervous system. It is unlikely that the numerous published microglia derivation protocols starting from human iPSCs would give rise to equivalent preparations, since the experimental conditions used by different methods vary considerably (eg, Muffat *et al*., 2016; Abud *et al*., 2017; Douvaras *et al*., 2017; Pandya *et al*., 2017; McQuade *et al*., 2018). We therefore sought to compare the gene expression signatures of human iPSC-derived microglia generated using different *in vitro* differentiation methods to begin to understand whether different methods have the potential to give rise to transcriptionally distinct microglia populations.

Two frequently used microglia derivation protocols, both of which achieve robust generation of microglia under serum-free conditions and in the presence of defined cytokines, were chosen. Specifically, human iPSCs were induced to undergo commitment to the myeloid lineage and differentiation into microglia following the methods by Douvaras and colleagues (Douvaras *et al*., 2017) or McQuade and coworkers (McQuade *et al*., 2018). Both of these methods have been used recently to model the involvement of human microglia in ALS pathophysiology (Allison *et al*., 2022; Kerk *et al*., 2022). In the context of the present study, we shall hereafter operationally refer to microglia generated using the Douvaras *et al* protocol as ‘Douvaras-microglia’, and microglia generated using the McQuade *et al* method as ‘McQuade-microglia’. Both protocols generated induced cultures highly enriched in cells expressing well-known microglial proteins such as IBA1 and P2RY12 (fractions of P2RY12^+^ cells: Douvaras-microglia, 96.3% ± 3.1%; McQuade-microglia, 95.2% ± 1.2%) (Fig. 1A, B). P2RY12 expression distinguishes microglia from peripheral macrophages (Butovsky *et al*., 2014). Induced cells displayed the morphologies of typical microglia (Fig. 1C).

**Figure 1.**
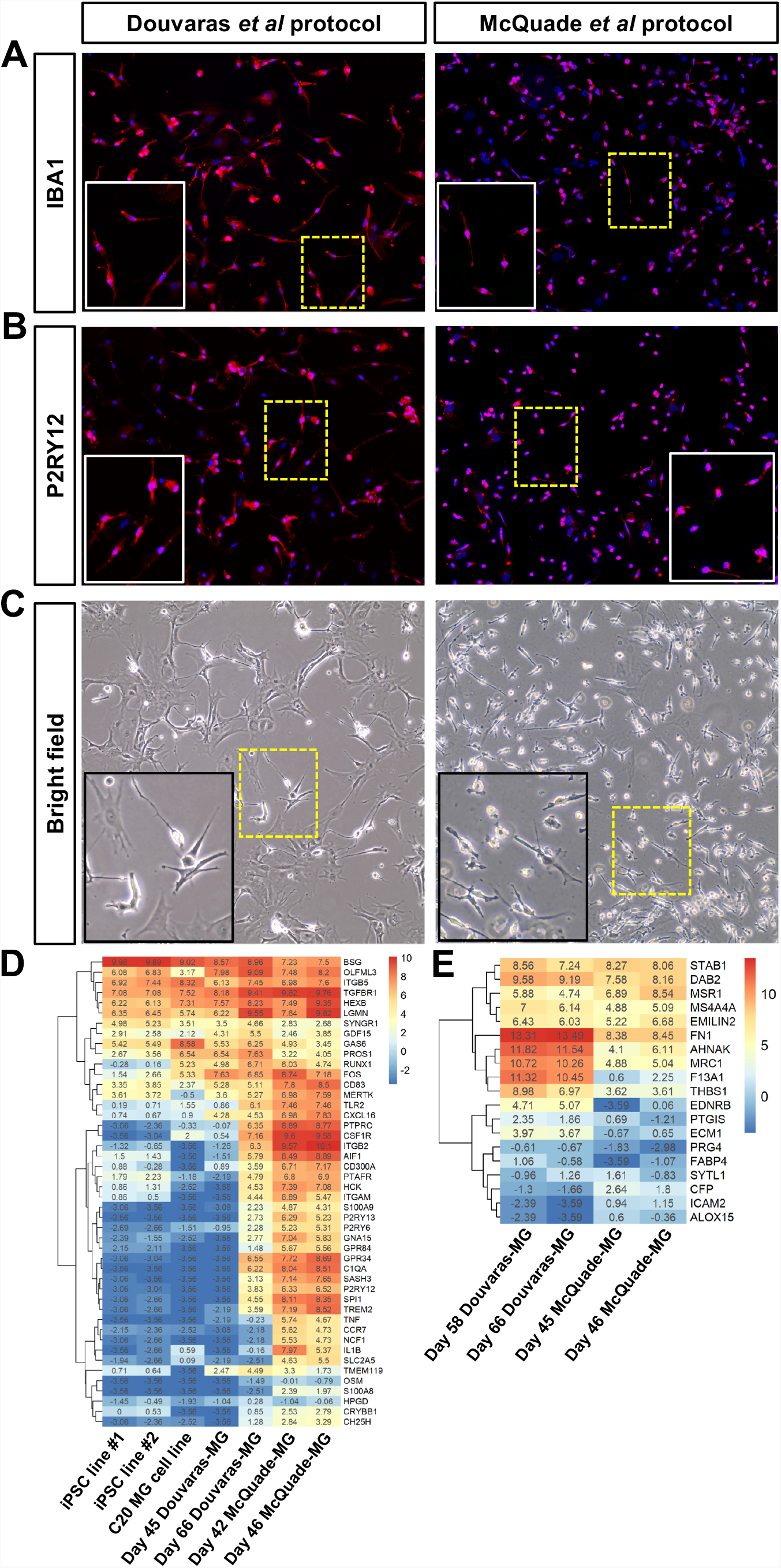
Characterization of human iPSC-derived microglia. **A, B)** Immunocytochemical analysis of either IBA1 (A) or P2RY12 (B) expression in microglia generated from iPSC line NRCM-1 examined at either differentiation day 48 (Douvaras *et al* protocol) or differentiation day 42 (McQuade *et al* protocol). Objective: 20X. Solid-line insets represent higher-magnification views of the areas delineated by dashed yellow boxes. **C)** Bright field views of microglia generated from iPSC line CS52iALS-C9n6.ISOxx examined at either differentiation day 51 (Douvaras *et al* protocol) or differentiation day 47 (McQuade *et al* protocol). Solid-line insets represent higher-magnification views of the areas delineated by dashed yellow boxes. **D)** Heatmap depicting the expression of known microglia genes in undifferentiated human iPSCs (two different lines from LINCS Data Portal open-source dataset LDS-1355), immortalized human microglia cell line C20 (from DEE2 repository open-source data), and human iPSC-derived microglia (‘MG’) generated in our lab with either the Douvaras or McQuade protocol and examined at the indicated differentiation days **E)** Heatmap depicting the expression of a macrophage-enriched gene signature in human iPSC-derived microglia (‘MG’) generated with either the Douvaras or McQuade protocol and examined at the indicated differentiation days. **D, E)** Cool colors indicate low expression and warm colors indicate increased/high expression, as shown in the color scale. The logCPM value (corresponding to the color scale) for each data point is listed in the heatmap. Subgroups of genes with correlated expression patterns are portrayed by the dendograms on the left side of the heatmaps.

To further characterize and compare both human iPSC-derived microglial cell preparations, we performed bulk RNAseq studies. Using transcriptomic data, we defined the expression level of a list of 49 microglial cell marker genes, compiled based on previous studies (Hickman *et al*., 2013; Butovsky *et al*., 2014; Zhang *et al*., 2014; Muffat *et al*., 2016), in Douvaras-microglia and McQuade-microglia. The majority of these 49 microglial genes were up-regulated in both Douvaras-microglia and McQuade-microglia when compared to undifferentiated human iPSCs and to the immortalized microglial cell line C20 (Garcia-Mesa *et al*., 2017) (Fig. 1D). The upregulated genes included *P2RY12*, in agreement with the immunocytochemical data shown in Figure 1B. In our hands, McQuade-microglia acquired a robust microglia gene-expression profile, compared to C20 microglia, 40–45 days after the start of the differentiation process. In comparison, Douvaras-microglia were transcriptionally similar to C20 microglia at *in vitro* differentiation day 45, and required an additional 15–20 days to exhibit a transcriptional signature similar to day-42–46 McQuade-microglia (Fig. 1D). Importantly, both microglia preparations displayed generally low expression of a previously described macrophage-specific gene signature (Hickman *et al*., 2013), except for a few genes, such as *DAB2, MSR1*, and *MRC1*, that are known to be expressed in both macrophages and microglia (Zeiner *et al*., 2019; Kobashi *et al*., 2020), which might point to the presence of common precursors of microglia and macrophages in those cultures (Fig. 1E). Together, these observations show that both microglial preparations display transcriptional signatures of physiological microglia, and suggest that McQuade-microglia undergo a more rapid developmental maturation *in vitro* than Douvaras-microglia.

To further examine the possibility that McQuade-microglia are more developmentally mature than Douvaras-microglia at comparable derivation time points, we next performed DEG analysis of RNAseq data collected from microglia generated with both methods. In order to minimize the possibility of detecting obvious differences resulting from different levels of *in vitro* developmental maturation, we compared cells displaying similar levels of expression of the 49 microglia markers listed in Figure 1D. Two separate Douvaras-microglia preparations (RNA collected at days *in vitro* 58 and 66) and two McQuade microglia preparations (RNA collected at days *in vitro* 45 and 46) were compared using an EdgeR analysis pipeline for DEG analysis (Chen *et al*., 2016). We detected 1949 genes that were significantly upregulated in McQuade-microglia compared to Douvaras-microglia, while 2867 genes were upregulated in the latter (adjusted p < 0.05 with no fold-change cutoff). *In silico* functional analyses (GO and KEGG analyses combined) of the DEGs between McQuade-microglia and Douvaras-microglia revealed that GO and KEGG terms involving biological functions typically expected from mature microglia (such as immune response, chemotaxis, motility, and migration) were over-represented in McQuade-microglia, suggesting that these cells were more developmentally mature than Douvaras-microglia (Fig. 2A). This analysis also showed that a number of biological processes/pathways suggestive of a more activated microglia phenotype were over-represented in McQuade-microglia, including “phagocytosis”, “apoptosis”, “NF-κB signaling pathway” (Fig. 2B).

**Figure 2.**
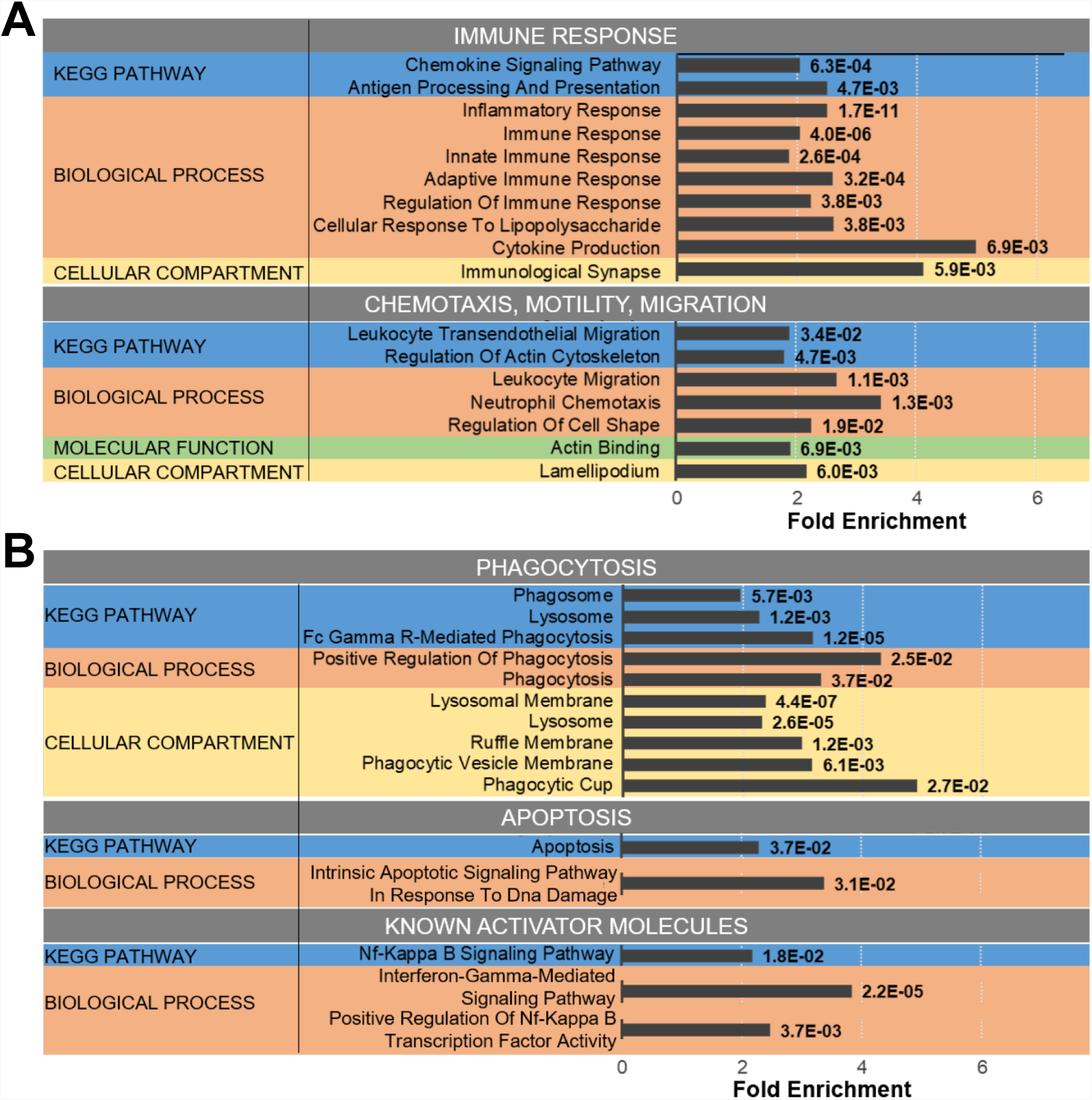
Human iPSC-derived microglia generated with different protocols exhibit distinct transcriptomic profiles. **A, B)** Functional analysis of DEGs between Douvaras-microglia (n=2; RNA collected at days *in vitro* 58 and 66) and McQuade-microglia (n=2; RNA collected at days *in vitro* 45 and 46). **B)** Barplot depicting gene ontology terms related to functions of more developmentally mature microglia (e.g., immune response, chemotaxis, motility, and migration) that are significantly over-represented in McQuade-microglia compared to Douvaras-microglia. The X-axis indicates “Fold Enrichment”, which is defined here and in succeeding figures as the enrichment of each term in the studied samples compared to its enrichment in the background population (number of DEGs in term/total number of DEGs) / (number of human genes in term/total number of genes in the human genome). The enrichment p-value is listed alongside its corresponding term. All functional categories with relevant terms are included, namely KEGG pathway, Biological Process, Molecular Function, and Cellular Compartment. Only non-disease-specific KEGG terms are included. **C)** Barplot depicting gene ontology terms associated with microglia activation (categorized under “phagocytosis”, “apoptosis”, and “known activator molecules”) that are significantly over-represented in McQuade-microglia compared to Douvaras-microglia. The X-axis indicates “Fold Enrichment” and the enrichment p-value is listed alongside its corresponding term. All functional categories with relevant terms are included, namely KEGG pathway, Biological Process, and Cellular Compartment. Only non-disease-specific KEGG terms are included.

Together, these observations provide evidence that McQuade-microglia have a transcriptional profile that suggests that they undergo a more rapid developmental maturation, and are more activated, than Douvaras-microglia across comparable *in vitro* differentiation trajectories.

### 3.2. Gene expression signatures of distinct in vivo microglia subtypes overlap with different human iPSC-derived microglia preparations

Based on the previous observations, we next investigated whether the transcriptomic divergence observed between Douvaras-microglia and McQuade-microglia might also be due to a different microglia-subtype composition in these preparations. The DEGs significantly upregulated in each preparation of microglia were compared to previously described gene signatures of regionally distinct microglia subtypes *in vivo*. At first, we compared the transcriptional profiles of Douvaras-microglia and McQuade-microglia to those of three distinct mouse microglia subtypes described by Grabert and colleagues (Grabert *et al*., 2016). These investigators isolated microglia from different regions of the adult mouse brain (cortex, hippocampus, striatum, and cerebellum) and identified three major subtypes based on their transcriptomic profiles. We observed that one of these three microglia subtypes was significantly enriched in Douvaras-microglia (Fisher’s exact test; p = 9×10^−4^), whereas a separate subtype was significantly enriched in McQuade-microglia (Fisher’s exact test; p = 0.018) (Fig. 3A). Importantly, the study by Grabert and coworkers (Grabert *et al*., 2016) had shown that the particular microglia gene signature that we observed to be enriched in Douvaras-microglia exhibited high expression in the mouse cerebellum and hippocampus, but low expression in the cortex and striatum. In contrast, the gene signature that we found to be enriched in McQuade microglia had reported high expression in the cortex and striatum, but low expression in the cerebellum and hippocampus (Grabert *et al*., 2016). These combined observations suggest that Douvaras-microglia and McQuade-microglia comprise regionally distinct microglial cell populations.

**Figure 3.**
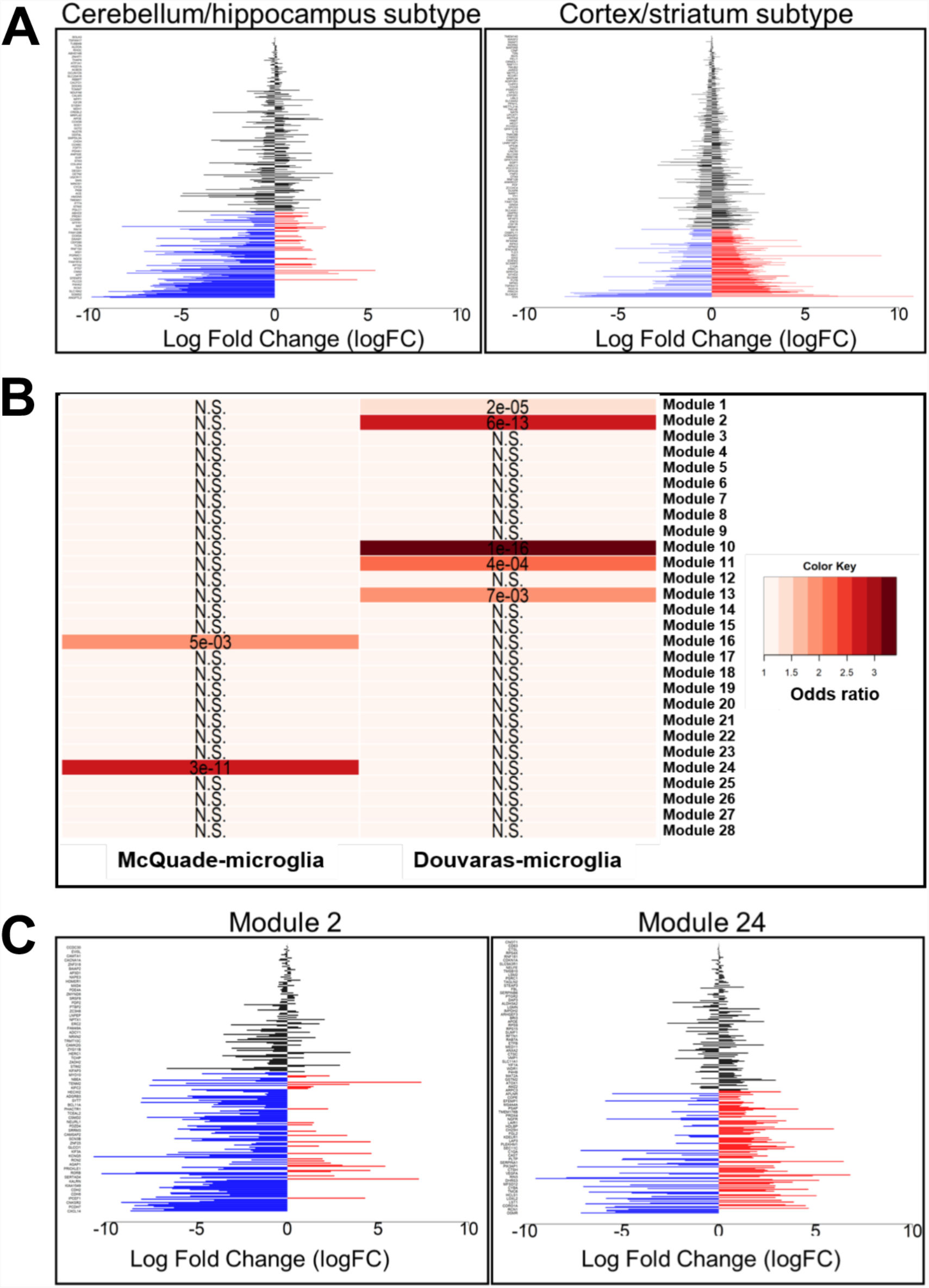
Overlap of regionally distinct gene signatures with iPSC-derived microglia generated using different methods. **A)** Barplots depicting the gene expression signatures of regionally distinct mouse microglia populations previously reported to preferentially localize to either the cerebellum and hippocampus or to the cortex and striatum (Grabert *et al*., 2016) in Douvaras-microglia and McQuade-microglia, respectively. Blue bars correspond to genes with significantly higher expression in Douvaras-microglia, whereas red bars correspond to genes with significantly higher expression in McQuade-microglia. Black bars are genes not significantly increased in microglia derived from either protocol. Log fold change in expression between the two protocols is displayed on the X-axis. Genes listed on the Y-axis can be found in Supplementary Table 1 or in the source publication (Grabert *et al*., 2016), where the “cortex/striatum subtype” and “cerebellum/hippocampus subtype” are respectively named “Cluster 1” and “Cluster 2”. **B)** Heatmap depicting the enrichment distribution of regionally distinct cell “modules” (Maniatis *et al*., 2019) in Douvaras-microglia and McQuade-microglia, by pairwise gene-overlap analysis. The color gradient represents the degree of association between the gene sets (odds ratio), and the overlapping p-value (Bonferroni-corrected) is listed in each cell. “N.S.” = not statistically significant. **C)** Barplots depicting the expression of Module 2 and Module 24 gene signatures in the microglia derived using the two different protocols. Blue bars are genes with significantly higher expression in Douvaras-microglia, whereas red bars are genes with significantly higher expression in McQuade-microglia. Black bars are genes not significantly increased in microglia derived using either protocol. Log fold change in expression between the two protocols is displayed on the X-axis. Genes listed on the Y-axis can be found in Supplementary Table 1, or in the source publication (Maniatis *et al*., 2019), where “Module 2” and “Module 24” are correspondingly named “human module 2” and “human module 24”.

To test this possibility further, we next cross-examined transcriptomic data from Douvaras-microglia and McQuade-microglia with previous results by Maniatis and colleagues, who conducted spatial transcriptomic analysis of *post-mortem* human spinal cord tissues and identified 28 regionally distinct cell “modules” (including but not restricted to microglia) (Maniatis *et al*., 2019). We observed that seven of these modules’ gene signatures significantly overlapped with upregulated genes in human iPSC-derived microglia generated using either tested protocol. Importantly, the five modules that were enriched in Douvaras-microglia did not overlap with the two modules we found enriched in McQuade-microglia (Fig. 3B). Based on the spatial analysis conducted by Maniatis and coworkers (Maniatis *et al*., 2019), we noticed that 3 of 5 modules enriched in Douvaras-microglia showed strong preferential expression for the gray matter of the human spinal cord, while the two modules enriched in McQuade-microglia were preferentially expressed in the white matter of the spinal cord. Two representative gene signatures enriched in Douvaras-microglia and McQuade-microglia, namely Module 2 (Fisher’s exact test; p-value for overlap with Douvaras-microglia = 6×10^−13^) and module 24 (Fisher’s exact test; p-value for overlap with McQuade-microglia = 3×10^−11^), are plotted in Figure 3C. These observations suggest further that microglia generated using the two *in vitro* differentiation methods under study are not equivalent, and raise the possibility that Douvaras-microglia and McQuade-microglia resemble gray-matter or white-matter microglia, respectively.

To examine the latter possibility further, gene signatures of both types of *in vitro* generated microglia were compared to a previously described transcriptomic characterization of white-matter microglia (Safaiyan *et al*., 2021). We observed that white-matter-associated genes displayed significantly increased expression in McQuade-microglia compared to Douvaras-microglia (Fisher’s exact test; overlapping p-value = 4.3×10^−3^) (Fig. 4A). In agreement with this finding, GO and KEGG analysis showed that, compared to Douvaras-microglia, McQuade-microglia displayed an over-representation of biological processes/pathways associated with white-matter microglia, such as “antigen processing and presentation”, “phagosome”, “chemokine signaling pathway”, and others (Lee *et al*., 2019; Amor *et al*., 2022) (Fig. 4B). In contrast, Douvaras-microglia exhibited over-represented functions characteristic of gray-matter microglia, such as interactions with astrocytes and neurons, and synaptic interactions (Fig. 4C).

**Figure 4.**
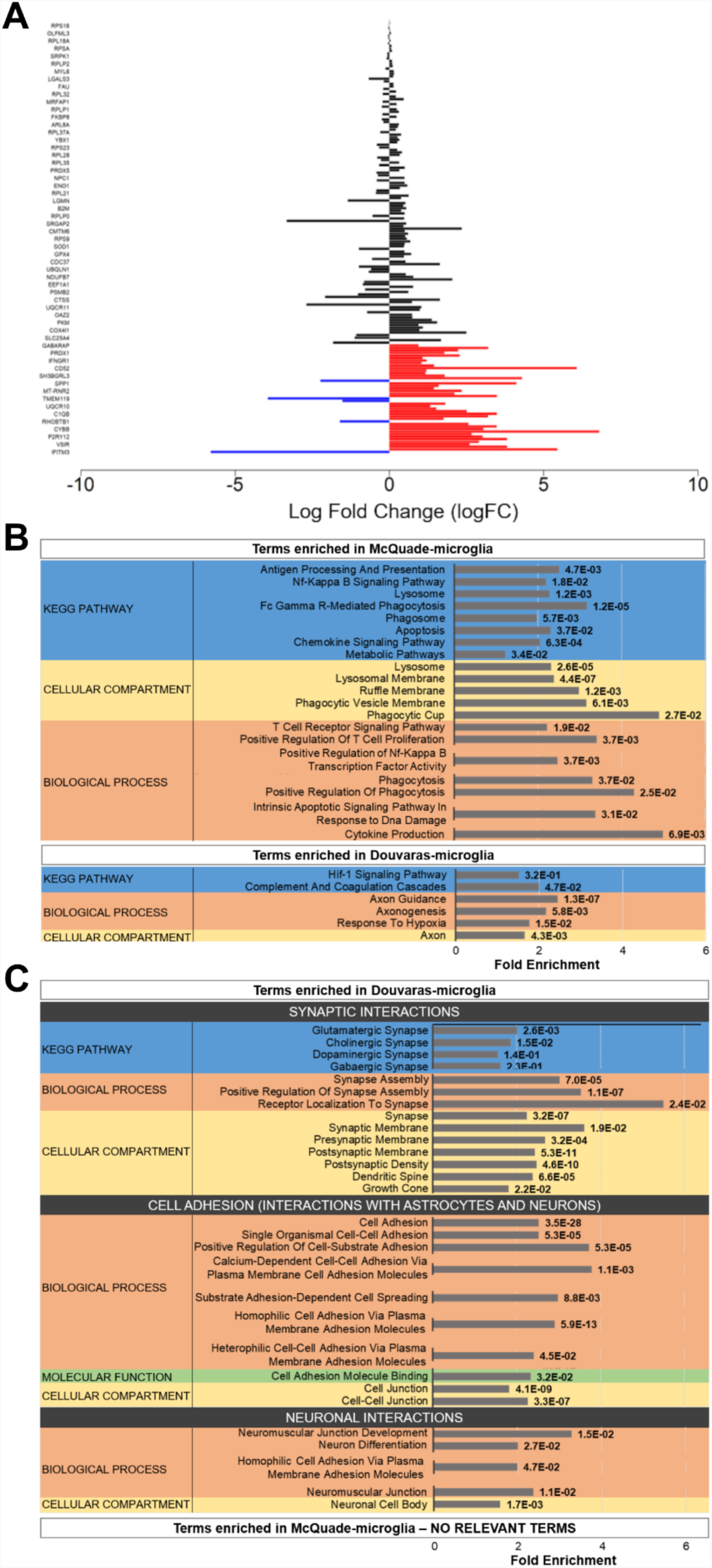
Gene signatures of iPSC-derived microglia generated with different protocols. **A)** Barplot depicting the expression of a white-matter-associated microglia (WAM) gene signature (Safaiyan *et al*., 2021) in microglia derived using the two methods under study. Blue bars correspond to genes with significantly increased expression in Douvaras-microglia, while red bars correspond to genes with significantly increased expression in McQuade-microglia. Black bars are genes not significantly increased in microglia derived from either protocol. Log fold change in expression between the two protocols is displayed on the X-axis. Genes listed on the Y-axis can be found in Supplementary Table S1 or in the source publication. **B)** Barplot depicting the gene ontology terms related to functions of white-matter microglia that are significantly over-represented in McQuade-microglia compared to Douvaras-microglia. The X-axis indicates “Fold Enrichment”. The enrichment p-value is listed alongside its corresponding term. All functional categories with relevant terms are included, namely KEGG pathway, Biological Process, and Cellular Compartment. Only non-disease-specific KEGG terms are included. **C)** Barplot depicting the gene ontology terms related to functions of gray-matter microglia (such as interactions with synapses, astrocytes, and neurons) that are significantly over-represented in Douvaras-microglia compared to McQuade-microglia. The X-axis indicates “Fold Enrichment”. The enrichment p-value is listed alongside its corresponding term. All functional categories with relevant terms are included, namely KEGG pathway, Biological Process, Molecular Function, and Cellular Compartment. Only non-disease-specific KEGG terms are included.

Taken together, these results provide evidence that the two different human iPSC-derived microglia preparations under study contain cell populations with distinct gene expression profiles correlating with regionally distinct microglial subpopulations identified *in vivo*. More specifically, Douvaras-microglia and McQuade-microglia exhibit gene signatures that overlap, at least in part, with microglia subpopulations present in the gray matter or white matter, respectively.

## 4. Discussion

The elucidation of the roles of microglia during the progression of neurodegenerative diseases, including ALS, has benefitted from the recent development of robust and reliable methods to generate microglia from human iPSCs, including iPSCs derived from patients affected by familial forms of these diseases and their matching genome-edited isogenic iPSCs. To-date, the few published studies using human iPSC-derived microglia to model ALS pathophysiology, which have used the two derivation methods investigated in the present study (Allison *et al*., 2022; Kerk *et al*., 2022), have not provided information on the subcellular composition of the microglia cultures that were examined. There is a growing need to gather more information on the diversity of different *in vitro* microglia experimental systems, stemming from the demonstrated spatial heterogeneity of microglia *in vivo* (Grabert *et al*., 2016; DePaula-Silva *et al*., 2019; Li *et al*., 2019; Masuda *et al*., 2019; Stratoulias *et al*., 2019; van der Poel *et al*., 2019; Masuda *et al*., 2020). Lack of this information can delay progress in the field of ALS research because it is likely that different subtypes of microglia interact with the variety of regionally distinct motor neurons that are affected by ALS in both the brain and spinal cord. Moreover, motor neuron degeneration in ALS is associated with pathological mechanisms in both the gray matter and white matter, which are populated by distinct microglia subtypes (Dachet *et al*., 2019; Maniatis *et al*., 2019; Cipollina *et al*., 2020). The future success of studies utilizing human iPSC-derived microglia to model neurodegenerative diseases will depend in large part on the ability to generate, and study, those specific microglial subtypes that are relevant to the pathobiological questions under investigation. In this work, we have provided evidence suggesting that by comparing, and integrating, different human iPSC-based microglia derivation protocols, it will be possible to gain insight into how to develop enhanced experimental systems that will take microglia heterogeneity, as well as other factors such as activation state, into consideration when modeling non-cell autonomous mechanisms of motor neuron degeneration in ALS using human iPSC-derived microglia.

Specific human iPSC-based neuronal or glial cell derivation methods are often chosen over other protocols because of length and easiness considerations. As important as these factors are, they carry the risk that the fastest or most amenable protocols may give rise to cultures enriched in only a few particular types of induced cells, which may be appropriate to study certain biological mechanisms but not others. The results of the present study suggest that it will be beneficial to the success of ALS-focused investigations utilizing human iPSC-derived microglia to conduct preliminary studies to select the most appropriate derivation methods on the basis of multiple criteria. The latter should include not only rapidity and easiness of the method, but also the cellular composition of the induced microglial cultures that would be ideal for the biological questions of interest (e.g., gray-matter versus white-matter microglia). The activation state of the induced microglia should also be considered, since intrinsically activated preparations may be less useful when studying dysregulated activation mechanisms in microglia derived from ALS patient iPSCs, compared to isogenic control cells.

The present work provides an example of studies that can be performed to address the above-mentioned questions, before certain (or combinations of) microglia derivation methods are selected. Specifically, our transcriptomic data suggest that human iPSC-derived microglia generated using the method described by McQuade and colleagues (McQuade *et al*., 2018) are enriched in cells resembling white-matter microglia. In contrast, microglial cultures generated as described by Douvaras and coworkers (Douvaras *et al*., 2017) are enriched in cells exhibiting gene signatures similar to those of gray-matter microglia. This finding provides support to the important notion that human iPSC-based approaches have the potential to give rise to distinct microglia subpopulations preferably suited to model specific mechanisms of ALS pathophysiology. We have also shown that microglia preparations generated using the McQuade method develop more rapidly and display more robust signs of activation than microglia derived using the Douvaras method at *in vitro* differentiation time points when these cells would usually be experimentally investigated. Thus, different derivation methods may offer both advantages and disadvantages in different contexts, further emphasizing the need to carefully consider the planned experimental strategies. It is reasonable to predict that when increased number of transcriptomic studies of different human iPSC-derived microglia become available, including the results of single-cell RNAseq, it will become increasingly recognized that 1) certain methods would be preferable for certain studies, and/or 2) a harmonised use of multiple methods might be the recommended approach to provide disease-relevant experimental systems, as this would generate a larger repertoire of microglia with optimal potential to reflect the heterogeneity present *in vivo*.

In conclusion, selecting the most informative human iPSC-based microglia derivation strategies is expected to become a key step in the development of disease-relevant experimental systems to model the involvement of microglia in multiple mechanisms of ALS pathophysiology. This goal will be facilitated by the growing understanding of microglia diversity *in vivo* and the mechanisms underlying the generation of this diversity. Combining this increased biological knowledge with a deeper characterization of the properties of *in vitro-*generated microglia will offer opportunities to optimize the development of advanced cellular systems to elucidate the specific roles of microglia in the pathophysiology of ALS, as well as other neurodegenerative diseases.

## Supplementary Materials

The following files are available online: Supplementary Table 1 and Supplementary Table 2.

## Authors contributions

Y.T. performed all tissue culture and microscopy studies. N.S.P. performed all bioinformatics studies. S.S. and N.S.P. wrote the manuscript. S.S. supervised the study. All authors have read and agreed to the published version of the manuscript.

## Acknowledgments

We thank Panagiotis Douvaras and Valentina Fossati for invaluable advice on microglia derivation, and Thomas Durcan and Jean-Francois Cloutier for providing access to scientific equipment. We also thank Rita Lo, Louise Thiry, Vincent Soubannier, Gilles Maussion, and Carol Chen for assistance, discussions, and advice.

## Data Sharing Statement

Read counts of the RNAseq data and the R code used for *in silico* analyses will be made available along with this publication on GitHub at https://github.com/nishapulimood/StifaniLab-iPSCmicroglia-protocol-comparison.

